# Deep conservation of histone variants in Thermococcales archaea

**DOI:** 10.1101/2021.09.07.455978

**Authors:** Kathryn M Stevens, Antoine Hocher, Tobias Warnecke

**Affiliations:** Medical Research Council London Institute of Medical Sciences, London, United Kingdom; Institute of Clinical Sciences, Faculty of Medicine, Imperial College London, London, United Kingdom

## Abstract

Histones are ubiquitous in eukaryotes where they assemble into nucleosomes, binding and wrapping DNA to form chromatin. One process to modify chromatin and regulate DNA accessibility is the replacement of histones in the nucleosome with paralogous variants. Histones are also present in archaea but whether and how histone variants contribute to the generation of different physiologically relevant chromatin states in these organisms remains largely unknown. Conservation of paralogs with distinct properties can provide *prima facie* evidence for defined functional roles. We recently revealed deep conservation of histone paralogs with different properties in the Methanobacteriales, but little is known experimentally about these histones. In contrast, the two histones of the model archaeon *Thermococcus kodakarensis*, HTkA and HTkB, have been examined in some depth, both *in vitro* and *in vivo*. HTkA and HTkB exhibit distinct DNA-binding behaviours and elicit unique transcriptional responses when deleted. Here, we consider the evolution of HTkA/B and their orthologs across the order Thermococcales. We find histones with signature HTkA- and HTkB-like properties to be present in almost all Thermococcales genomes. Phylogenetic analysis indicates the presence of one HTkA- and one HTkB-like histone in the ancestor of Thermococcales and long-term maintenance of these two paralogs throughout Thermococcales diversification. Our results support the notion that archaea and eukaryotes have convergently evolved histone variants that carry out distinct adaptive functions. Intriguingly, we also detect more highly diverged histone-fold proteins, related to those found in some bacteria, in several Thermococcales genomes. The functions of these bacteria-type histones remain entirely unknown, but structural modelling suggests that they can form heterodimers with HTkA/B-like histones.

## Introduction

The ability of eukaryotic cells to respond to environmental change and regulate transcription relies on dynamic control of DNA accessibility through chromatin alterations. This involves many different processes, including the addition/removal of histone modifications and the exchange of histone proteins for paralogous variants. Such variants can modify structural properties of the nucleosome or change how it interacts with its binding partners (1–3). For example, macroH2A has a large C-terminal domain and precipitates transcriptional repression (2, 4) while cenH3, a fast-evolving H3 variant, is specifically localised to centromeres and involved in chromosome segregation (1, 5). Importantly, significant functional changes can come from small differences in sequence. H3.3, for example, is deposited in a replication-independent manner in actively transcribed regions of the genome (1) and important for mammalian development (6, 7), but differs from its paralog H3.1 by only five amino acid.

Histones are not exclusive to eukaryotes. Archaeal histone proteins, first discovered in *Methanothermus fervidus* (8, 9), have since been identified in diverse archaeal lineages (10, 11) and are often highly expressed (11). Eukaryotic and archaeal histones share a conserved histone fold (HF) domain, form dimers and tetramers that are structurally very similar, and bind DNA non-specifically, albeit with a preference for more bendable sequences (12–16). Unlike eukaryotic histones, almost all archaeal histones lack long terminal extensions (“tails”) (10) but have the capacity to form homo- as well as heterodimers and to assemble into long oligomeric structures (16). These extended, flexible complexes (17) can, in theory, consist of different histone paralogs, providing opportunities for chromatin state modulation through the exchange of histones with different properties (18). In fact, many archaea encode two or more sequence-divergent histone paralogs, but whether these paralogs have defined functional roles akin to eukaryotic histone variants, and whether their expression and assembly change dynamically to mediate adaptive chromatin states, remains poorly understood.

What we do know from prior experimental work is that archaeal histone paralogs are more than mere copy number variants. The two histones of *M. fervidus* (HMfA, HMfB), for example, display differences in DNA binding affinity (19). Compared to HMfA, recombinant HMfB also induces more positive supercoiling upon binding to plasmid DNA and forms a more compact complex as inferred from gel-shift and tethered particle motion experiments (20, 21). There are also differences between HMfA and HMfB in their relative expression during the growth cycle: in early exponential phase, HMfA is more highly expressed than HMfB, whose expression level increases towards stationary phase to reach an almost equal ratio between the two (20). The different properties of *M. fervidus* histones are consistent with the hypothesis that the two paralogs may have distinct functions in nucleoid biology, but whether the properties are physiologically relevant and affect organismal fitness has rarely been addressed explicitly.

Recently, we considered this question using an evolutionary approach. We identified histone paralogs in the order Methanobacteriales that exhibit distinct structural properties and have been maintained over hundreds of millions of years (18), indicative of the importance of each individual paralog for fitness. Structural modelling identified histone variants that prevent stable tetramerization and might act as *capstones* that limit further extension when incorporated into a histone oligomer, providing a potential pathway to dynamically alter chromatin state. Are the Methanobacteriales unique or are there other clades of archaea with histone paralogs that have been maintained over long periods of time? And do these paralogs also show conserved and distinct structural properties?

Here, we consider archaea in the order Thermococcales, which includes the model archaea *Pyrococcus furiosus* and *Pyrococcus abyssi* as well as *T. kodakarensis*, which has served as a model species for the *in vivo* study of archaeal histones. Thanks to the efforts of Santangelo and co-workers in particular, its two histones – HTkA (TK1413) and HTkB (TK2289) – are arguably the best characterized paralogs *in vivo*. Similar to HMfA and HMfB in *M. fervidus*, HTkA and HTkB can assemble into long oligomers both *in vitro* and *in vivo* (16, 17, 22, 23). The two histones differ from one another at 11 out of 67 residues (84% identity) and have several distinct properties. HTkA is the more highly expressed paralog, at least in exponential phase, where it makes up 1.1% of the proteome compared to 0.66% for HTkB (11). Together, they are abundant enough to coat the entire *T. kodakarensis* genome (24). HTkB has been shown to bind to DNA more strongly than HTkA and to form more compact complexes, which show faster migration during agarose gel electrophoresis (25). Deletion of each histone individually results in overlapping but distinct perturbations of the transcriptome (22, 26). Notably, HTkB-deficient cells exhibit reduced growth, possibly due to changes in the expression, not seen in strains lacking HTkA, of genes that encode translation factors and ribosomal proteins (26). Deletion of *htkA* but not *htkB* leads to downregulation of hypothetical membrane proteins and prevents transformation of *T. kodakarensis*, suggesting HTkA alone plays a critical role in DNA uptake and/or integration (26).

In this study, we show that histone paralogs with HTkA- and HTkB-like properties are present across the Thermococcales, including Thermococcus, Pyrococcus, and Palaeococus *spp*.. We use structural modelling to show that, in most Thermococcales, HTkB-like histones are predicted to exhibit stronger DNA binding than those with HTkA-like properties. Phylogenetic analysis reveals that HTkA-like histones share a common ancestor to the exclusion of HTkB-like histones and *vice versa*, suggesting that the last common ancestor of the Thermococcales already encoded an HTkA-like and an HTkB-like histone, each of which has been maintained throughout the diversification of this clade for (very) approximately 750 million years (27). The long-term preservation of these two paralogs across the order Thermococcales supports the notion that HTkA/B in *T. kodakarensis* (and their orthologs in other Thermococcales) make unique contributions to genome function and fitness. These findings add further evidence that histone variants exist in archaea, evolving in parallel to those in eukaryotes.

Intriguingly, many Thermococcales archaea encode additional types of histone-fold proteins that are similar to histone-fold proteins found in some bacteria (28). One of these consists of an end-to-end duplication of the histone fold and is rarely found in archaea outside the Thermococcales. Within the Thermococcales, these bacteria-type histones are – while widespread – frequently lost, in contrast to HTkA/B. Their physiological roles remain unknown, but structural modelling suggest that they are able to heterodimerize with HTkA/B and might therefore further diversify histone-based chromatin states in these archaea.

## Results and Discussion

### Almost all Thermococcales have histone paralogs with HTkA- and HTkB-like properties

To identify putative histone proteins across the Thermococcales, we scanned 61 predicted proteomes using HMM models and iterative jackhmmer searches (see Methods). Histones with a single histone-fold domain (similar to canonical archaeal HMf-like histones) were found in all genomes (Fig 1c, Table S1). We also recovered putative histone-fold proteins similar to those found in some bacteria (28), which we discuss further below. A principal component analysis of the HMf-like archaeal histones based on their amino acid properties and isoelectric points (see Methods) suggests that histones can be assigned to one of two groups. One of these groups contains HTkA, the other HTkB (Fig 1a). This is consistent with a previous classification effort that also recovered two major groups of Thermococcales histones (29). Amino acid identities at several residues along the histone fold differ systematically between groups and are diagnostic of group membership. For example, tyrosine is always found at position 35 (Y35) in HTkA-like histones whereas HTkB-like histones have a positively charged lysine (K, 60 out of 62) or histidine (H, 2 out of 62). Similarly, glutamic acid at position 18 (E18) is present in 59 out of the 61 HTkA-like but none of the HTkB-like histones (Fig 1b).

**Figure 1.**
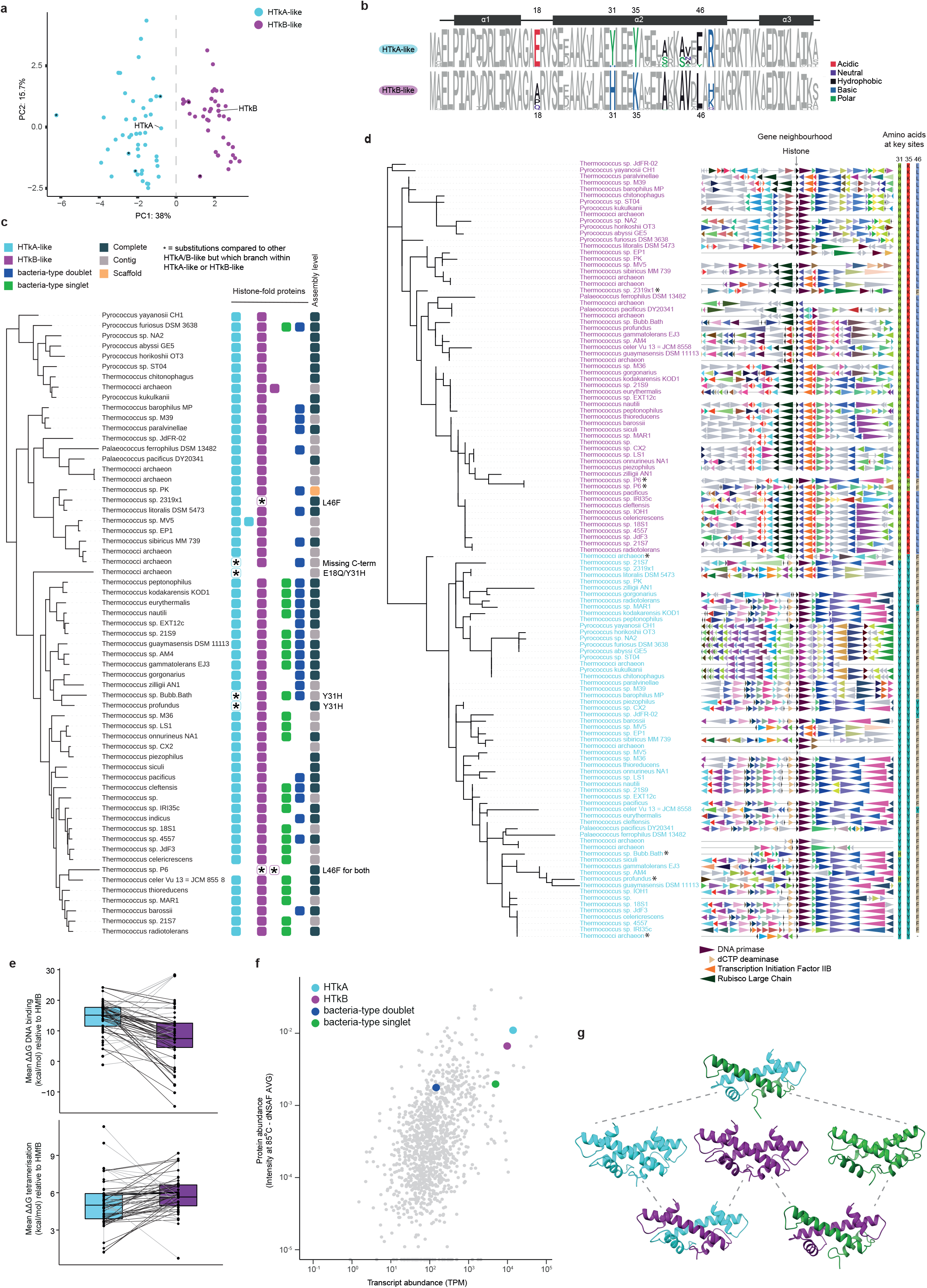
**a.** Principle component analysis of Thermococcales HMf-like histones based on amino acid properties (AAStats, see Methods). Histones that cluster with either HTkA or HTkB along the first principal component are coloured accordingly (HTkA-like histones in blue, HTkB-like in purple). Histones with substitutions at discriminative positions (see panel b) are highlighted with an asterisk. **b.** Sequence logos showing amino acid composition of HTkA- and HTkB-like histones across 61 Thermococcales. Amino acids that differ noticeably between the two groups are coloured. Positions are numbered relative to HMfB from *Methanothermus fervidus* to facilitate comparison with prior studies. Residues with known functional importance in archaea from previous experiments are highlighted on the secondary structure cartoon as in (16) **c.** Protein-level phylogenetic tree of all HMf-like Thermococcales histones. Genes in the neighbourhood are coloured to indicate ortholog identity (see Methods). Note that, while the gene neighbourhood is broadly conserved for HTkA-vis-à-vis HTkB-like orthologs, some HTkB-like genes (e.g. in *Thermococcus chitonophagus*) have a 3’ neighbourhood normally found for HTkA. This is owing to a large genomic rearrangement event in which HTkB served as the breakpoint (not shown). **d.** GTDB species tree for all Thermococcales in the dataset indicating presence/absence of histones of a particular type in each genome. Each square represents one histone and is coloured by histone type. **e.** Predicted DNA binding affinity (top) and tetramer stability (bottom) for HTkA/B-like paralogs. Lines connect paralogs from the same genome. **f.** Protein and transcript abundance (see Methods) of genes in *T. kodakarensis.* TPM: transcripts per million. **g.** AlphaFold-predicted homodimeric structures of TK1040 (green), HTkA (light blue) and HTkB (purple) and heterodimers of HTkA and HTkB (light blue/purple), HTkA and TK1040 (light blue/green), and HTkB and TK1040 (purple/green).

For some of these residues, we know from prior *in vitro* studies – as well as structural modelling – that amino acid identity can affect specific histone properties (18, 30). For example, substituting Y for K at residue 35 (the amino acids seen in HTKA- and HTKB-like histones, respectively), increases the stability of recombinant histone HFoB from *Methanobacterium formicicum* (31). In addition, evidence from mass spectrometry indicates that K35 in HTkB from *T. kodakarensis* and *Thermococcus gammatolerans* is acetylated *in vivo* (32) although stoichiometry and functional significance of this modification remain to be determined. A tyrosine at the same position in HTkA removes the potential for acetylation. E18 in HMfB forms an intermolecular salt bridge with K53, which helps to stabilise the interaction between monomers in the histone dimer (15, 33). Mutating E18 to proline does not alter DNA binding (30) but loss of the intermonomer salt bridge may result in less rigid dimer structures. Finally, having leucine (L) or phenylalanine (F) at residue 46 has no obvious effect on DNA binding in HMfA/B, but the residue, located at the interface between dimers, is important for tetramer formation (30, 34).

We considered how amino acid differences between HTkA- and HTkB-like histones affect two key aspects of the histone-DNA complex: DNA affinity and tetramerization strength, a proxy for tetramer stability. Using a structural modelling approach, we find that predicted DNA binding for HTkB-like paralogs is, in most cases, stronger than for HTkA-like paralogs (Fig 1e, see Methods). This is in line with experimental findings that HTkB binds to DNA more tightly and forms a more compact complex with DNA than HTkA (25). In contrast, predicted tetramerization strength does not strongly discriminate HTkA- from HTkB-like histones.

### HtkA- and HTkB-like histones form ancient paralogous groups

Almost all Thermococcales have both an HTkA-like and an HTkB-like histone (Fig 1c). This is consistent with (but not sufficient to demonstrate) ancient paralogy. To unravel the evolutionary history of Thermoccoccales histones, we used RaxML-NG (35) to build phylogenetic trees of all 123 canonical histones found across the 61 genomes in our analysis (see Methods). We find that HTkA-like and HTkB-like paralogs neatly separate into two groups defined by their position on the tree (Fig 1d). This pattern of separation indicates that one HTkA- and one HTkB-like histone were present in the last common ancestor of Thermococcales. The observation that both paralogs have been maintained along divergent Thermococcales lineages strongly suggests that at least some of the amino acid differences between them are functionally important and under selection, notwithstanding the fact that *T. kodakarensis* cells can tolerate the deletion of either histone under standard culture conditions (26). Along with our recent report of ancient histone paralogs in the Methanobacteriales (18), this finding provides further evidence that histone variants exist in archaea, evolving in parallel to those in eukaryotes.

### Some Thermococcales encode histone-fold proteins similar to those found in bacteria

Our survey also revealed that, alongside the HTkA/B-like histones, many Thermococcales genomes encode histone-fold proteins similar to those found in some bacteria (Fig 1c, Table S1), which harbour either a single or two (pseudodimeric) histone fold domains (28). We will refer to these as bacteria-type singlets and doublets, respectively. In contrast to HTkA/B-like histones their distribution across the Thermococcales is noticeably patchier and neither type is present in the closest sister clades (Methanofastidiosa, Theionarchaea). Looking more widely across archaea, the bacteria-type doublets are additionally only found in Methanocaldococcus and Archaeoglobus species (Hocher *et al.* in preparation), which are also hyperthermophiles, suggesting that the last common ancestor of the Thermococcales likely acquired this gene via horizontal transfer. These predicted histone-fold proteins have only recently been recognized and await functional characterization. The only functional data we have at present comes from transcriptome/proteome profiling: the relative expression levels of these genes in *T. kodakarensis* (singlet: TK1040; doublet: TK0750) are lower than those of HTkA/B-like histones at both the transcript and protein level (Fig 1f) (36, 37). Together, they make up 0.37% of the measured exponential-phase proteome compared to 0.66% for HTkB and 1.1% for HTkA. TK1040 was previously identified in *T. kodakarensis* chromatin fractions (38), suggesting a (direct or indirect) association with DNA. However, the same study estimated that less than 1% of the amount of chromatin-associated proteins were attributable to TK1040. We therefore consider it unlikely that these histones are global organizers of DNA similar to HTkA/B-like histones, but might modulate chromatin state, either locally or globally, in response to environmental change. In both *P. furiosus* and *T. kodakarensis*, the bacterial-type doublets are under the control of the heat shock regulator Phr (encoded by PF1790 and TK2291, respectively) and upregulated upon Phr deletion, suggesting a potential role in response to heat shock in these archaea (39, 40). The *T. kodakarensis* doublet is also downregulated at lower temperatures, similar to HTkA/B (11), further consistent with a role in temperature adaptation.

Can these histone fold proteins interact with HTkA/B-like histones? We used AlphaFold (41) to predict the structure of combinations of HTkA, HTkB and the bacterial HF singlet from *T. kodakarensis*. Using this approach, all three are predicted to form homodimers and, as expected, HTkA and HTkB form a stable heterodimer. When a combination of either HTkA or HTkB and the bacterial singlet are used, Alphafold also predicts that these will form a heterodimer (Fig 1g). The presence of non-HMf-like HF proteins in Thermococcales genomes adds to the potential functional diversity of histone-based chromatin in these species and may dynamically alter DNA accessibility at different stages of cell growth or in response to environmental challenges. Further experimental investigation is required, however, before meaningful conclusions can be drawn in this regard, including whether they do interact, both structurally and functionally, with the canonical HTkA/B-like histones.

## Methods

### Identification of histones in Thermococcales genomes

Protein sets, genomes and GFF files for all available genomes of class Thermococci were downloaded from GenBank (https://www.ncbi.nlm.nih.gov/assembly) using taxid 183968 [accessed on 2021-05-27]. Genomes not present in the GTDB tree (https://gtdb.ecogenomic.org, accessed 2021-08-01, see below) were removed. Two species which were annotated as Thermococci in NCBI but branched outside the main group on the GTDB tree were removed from the analysis, leaving a final set of 61 genomes, all from the order Thermococcales. Protein sequences were predicted using Prodigal v2.6.3 (42) where not provided through GenBank. Histone proteins were extracted from the protein sets through HMM searches using HMMER v3.3.1 (hmmsearch --noali) (43, 44) using Pfam models CBFD_NFYB_HMF and DUF1931 (45) as well as a Jackhmmer searches using the singlet and doublet histones from bacteria as a seed used by others (28). Some proteins incorrectly identified as histones at this stage were manually filtered out.

### Classification of Thermococcales histones into HtkA-like and HtkB-like groups

HMf-like histones were aligned using MAFFT (46) (–localpair --maxiterate 1000). Histones were clustered based on amino acid composition of their peptide sequences using AAStats from the R package Seqinr (47). Histones which clustered with HTkA and HTkB were assigned HTkA- or HTkB-like status, respectively (see Fig 1a). 20 maximum likelihood phylogenetic trees were built using Raxml-NG (35) with the LG+G4 model of evolution as suggested by ModelTest-NG (48). The unrooted best maximum likelihood tree is shown. All trees were plotted using iTOL (49). Orthologous genes in the genomic neighbourhood of each histone were highlighted on the tree using Genespy as best reciprocal hits (50). Orthologs were identified by performing reciprocal best hits for each genome against *T. kodakarensis* using BLAST (51), retaining those which have a similarity score of >40% and are within 20% length of one another (52). To generate sequence logos, histones were aligned using MAFFT (46) (–localpair --maxiterate 1000) and visualized using ggseqlogo in R (53).

### Predicted DNA binding and tetramerisation

Predicted DNA-binding and interaction (tetramerization) strength between dimers for Thermococcales species was computed as in (18). In brief, sequences were aligned to HMfB and substitutions were mapped onto a tetrameric model of HMfB (extracted from PDB structure 5t5k) using FoldX (54) to generate models of homotetramers with DNA. Structures were then energy-minimised using AmberTools (55) and binding affinity was calculated using an MMPBSA approach with the ff14SB forcefield (56). ΔΔG was calculated relative to HMfB. The mean value for five replicates is shown for each model.

### Species tree

The archaeal species tree was downloaded from GTDB (https://gtdb.ecogenomic.org) on 2021-08-01.

### Expression data

Expression data for *T. kodakarensis* was obtained from primary sources and NCBI’s Gene Expression Omnibus (GEO) (57). Proteomics data from (36) was processed as in (11). Protein abundance at 85°C is shown. Transcript abundance data was obtained from (37) and is shown as transcripts per million (TPM).

### Bacterial histone fold singlet and HTkA/B structure prediction

The AlphaFold v2.0 (41) collab notebook (https://colab.research.google.com/github/deepmind/alphafold/blob/main/notebooks/AlphaFold.ipynb <u>accessed 18-08-21</u>) was used to predict the structure of HTkA, HTkB and the bacterial singlet HF protein (TK1040) as homodimers and all heterodimer combinations. The MSA method used was jackhmmer, and models were ranked by PTMscore. The top model is shown for all homodimers and heterodimers. Images shown were generated using UCSF, ChimeraX developed by the Resource for Biocomputing, Visualization, and Informatics at the University of California, San Francisco, with support from National Institutes of Health (R01-GM129325) and the Office of Cyber Infrastructure and Computational Biology, National Institute of Allergy and Infectious Diseases (58).

